# High Resolution Acoustic Mapping of Gelatin-Based Soft Tissue Phantoms

**DOI:** 10.1101/2023.05.10.540075

**Authors:** Heba M. Badawe, Petra Raad, Massoud L. Khraiche

## Abstract

**Background:** Utilizing spatially and temporally uniform tissue-mimicking phantoms for ultrasonic applications can facilitate the characterization of beam distortion and attenuation. The implementation of acoustic phantoms can enhance the efficacy of ultrasound therapy or imaging by providing guidance on optimal ultrasonic parameters, such as frequency and power. The efficacy of phantoms is heavily dependent on the accuracy and reliability of measurement techniques employed for assessing their acoustic properties.

**Purpose:** The work aims to develop, build, and characterize, via high resolution acoustic mapping, Gelatin-Based ultrasound (US) soft tissue phantoms. To that effect, we built acoustic maps of the intensity distribution of US waves passing through the phantoms and studied the effect of gelatin concentrations and US frequency, duty cycle, and applied voltage on the acoustic intensity and focal region of the US waves. The methodology developed here offers well characterized and reproducible Gelatin-Based US phantoms for soft tissue (both acoustically and mechanically).

**Methods:** We developed gelatin-based phantoms, with conveniently adjustable parameters and measured, with high resolution, the acoustic attenuation of ultrasound waves when encountering the gelatin phantoms. This was done via a motorized acoustic system built for 3D-acoustic mapping of ultrasound waves. Mechanical assessment of the phantoms’ elasticity was carried out through unconfined compression tests. We characterized tissue mimicking phantoms with realistic acoustic properties and mechanical elasticity, emphasizing the effect of varying gelatin concentration on the ultrasound maximal intensity, thus causing acoustic attenuation throughout the acoustic profile. For validation, we used computational simulations to compare our data to predicted acoustical outcomes.

**Results:** Our results show high-resolution mapping of US waves in fluid with and without Gelatin-Based phantoms. We also confirm the impact of recipe and gelatin concentration on mechanical and acoustic characterization of phantoms. The density of the gelatin-based phantoms scales with the Young’s modulus. When characterizing the acoustic profiles of the different ultrasound transducers, the focal areas increased systematically as a function of increasing applied voltage and duty cycle yet decreased as a function of increased ultrasonic frequency.

**Conclusions:** We developed a Gelatin-Based US phantoms are a reliable and reproduce tool for examining the acoustic attenuations taking place as a function of increased tissue elasticity and stiffness. High resolution acoustic maps of the intensity distribution of US can provide essential information on the spatial changes in US wave intensity and focal point enabling a more in-depth examination of the effect of tissue on US waves.

## Introduction

Ultrasound is a non-invasive modality used clinically for diagnostic and therapeutic purposes. [1]. Ultrasonic energy beams can be focused at target tissues with high spatial and temporal resolution overcoming, to a certain extent, tissue inhomogeneity [2]. Ultrasound has a wide range of applications, including both diagnosis and treatment, the latter depending on the intensity and frequency used. For instance, low-intensity focused ultrasound (LIFUS) has demonstrated neuromodulation capabilities, enabling the reversible excitation or inhibition of neurons with no reported brain damage. [3-9], whereas high intensity focused ultrasound (HIFU) causes lesions and ablates cancerous tumors irreversibly [4, 10], depending on exposure time and tissue temperature elevation, thus becoming a potential neurosurgery technique. Besides neuromodulation, ultrasound is mainly used in image-guided tissue visualization in pre-clinical and clinical applications, allowing treatment planning and online assessment of ultrasound interactions with target tissues [11]. Even though ultrasonic imaging has some limitations in the overall showcasing of tissue anatomy, it provides valuable information on tissue dynamics, blood flow, and soft tissues stiffness and elasticity [12].

Applying ultrasound waves to spatially and temporally uniform tissue-mimicking phantoms can aid in understanding the amount of beam distortion and attenuation taking place [13]. Constructing a phantom that mimics the acoustic, optical, electrical, thermal, and mechanical properties of biological tissues can be crucial for preclinical and clinical applications of US [14, 15]. A tissue phantom can be homogeneous or heterogeneous, mimicking the various layers of tissue in question. Yuan et al. constructed a phantom representing the human thigh with an embedded tumor enclosing the fat, muscles, and bones [16]. Li et al. formulated a composition of different oils and chemical materials to represent the breasts and surrounding tissues in an attempt to study the interactions between such materials with ultrasound and microwave radiations [17]. T Tuning the various tissue properties is feasible during the design and fabrication stage through different chemical additives and mechanical compositions to account for the different geometrical shapes of target tissues [18]. Even for medical practices and surgical purposes, silicone-based phantoms were constructed to assess the flexibility, accuracy, and functionality of medical tools [19]. In the thermal ablation field, ultrasound or magnetic resonance imaging modalities can be guided by the tumor phantom inserted within the normal tissue phantom for targeted ablation based on tumor color change at higher temperatures [20, 21]. Takagi et al. developed a color-coded, thermochromic liquid-crystal tissue-mimicking phantom, composed of two layers with different temperature sensitivity ranges, for better visualization of the distribution of high-intensity focused ultrasound waves around the focal point [22]. High-intensity ultrasound attenuation can also be detected based on the tissue displacement induced within the phantom in the direction of ultrasound beams, and can be visualized through ultrasound imaging [23].

In this work, we focus on building high-resolution acoustic maps of ultrasound waves propagating in water, while using maps to study the effect of ultrasound frequency, duty cycle, and voltage on the maximal acoustic intensity and area of the focal region. We then develop gelatin-based phantoms, with three different gelatin concentrations that represent various soft tissues with similar mechanical and acoustic characteristic, to account for the attenuation in the maximal acoustic intensity. After that, we study the effect of increasing gelatin concentration on the compressive stress and strain and the mean Young’s modulus of the gelatin-based phantoms. Increasing the gelatin concentration rendered the phantom molds firmer and denser, thus recording higher Young’s moduli and causing more acoustic attenuation with greater power loss per unit area. The performed tests include compression loading with a sensitive load-cell installed for mechanical testing, while acoustic measurements were performed based on the acquired voltage signals from the hydrophone inserted in a water bath in front of the face of the ultrasound transducer. High-resolution intensity maps were obtained to illustrate the spread of ultrasound waves across the acoustic profile, for visual inspection of the intensity drop and increased acoustic attenuation with an increase in gelatin concentration. Simulation studies were also performed to validate this change for more precise visualization of the ultrasound beam distortion.

## Materials and Methods

### Experimental Setup

A motorized system was assembled as in Figure 1 to map the acoustic profile of the ultrasound transducer at 30 µm resolution and generate heat maps that illustrate the transfer of intensity of ultrasound waves in a specific volume of water. This helped characterize the acoustic intensity at the focal region while using three different drive voltages of 110 mV, 120 mV and 165 mV, with transducers with three different frequencies of 1 MHz, 5 MHz, and 7.5 MHz. Following that, and to characterize the effect of gelatin-based phantoms with three different gelatin concentrations on the acoustic intensity and attenuation, horizontal planes facing the transducer were scanned; the gelatin-based phantoms were positioned between the transducer probe and hydrophone (Figure 2).

**Figure 1.**
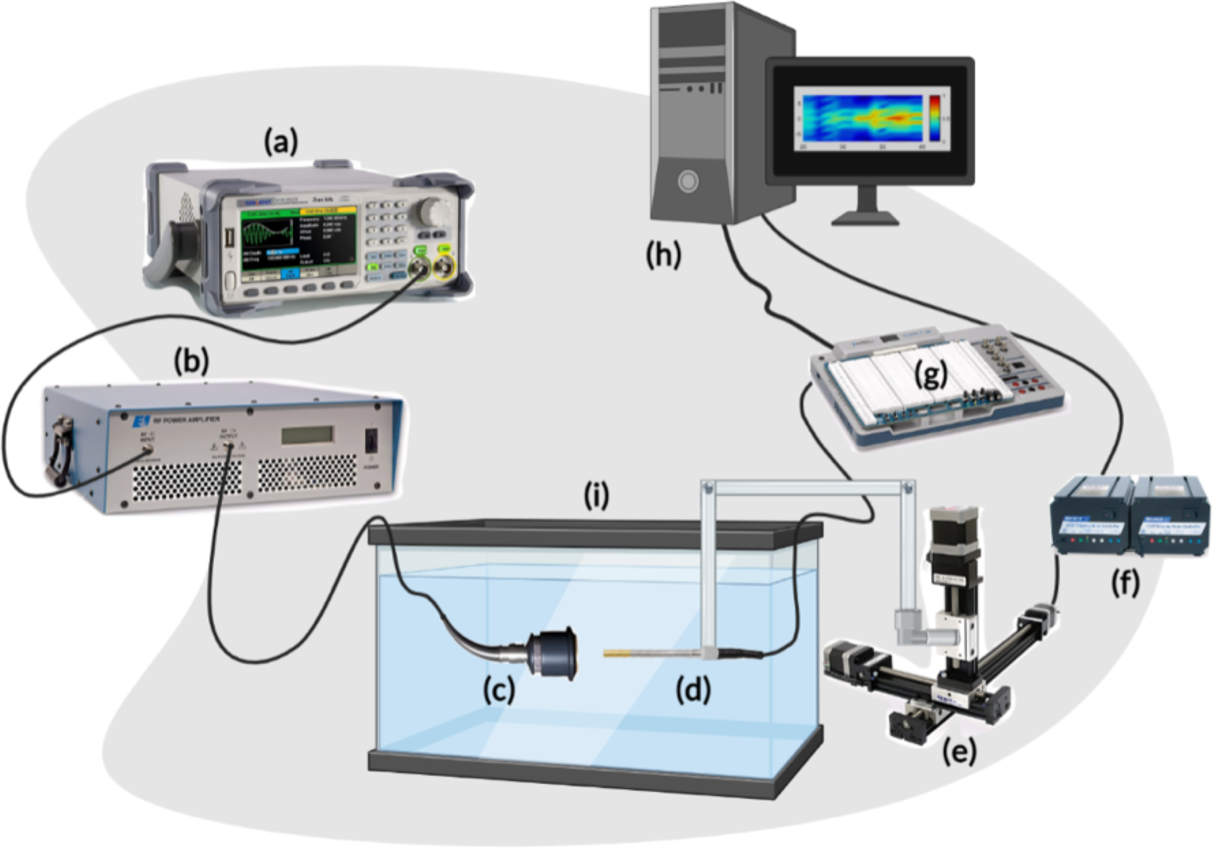
Scheme of the acoustic profile set-up. (a) represents the function generator producing a pulsed sine wave. (b) is the RF power amplifier which amplifies the sinusoidal RF signal that is converted to ultrasonic pressure waves by the ultrasound transducer (c). The hydrophone (d) is driven by the 3-D motorized system I, powered by the VXM stepping motor controller (f), and converts the mechanical signals received from the transducer into electrical signals while scanning a 3D volume in front of the face of the transducer. The voltage signals from the hydrophone are received by the Elvis III data acquisition board (DAQ) (g) and relayed back to the PC (h) for further processing. Both the transducer and hydrophone are submerged in the degassed water tank (i) of dimensions 40 cm x 20 cm x 16 cm.

**Figure 2.**
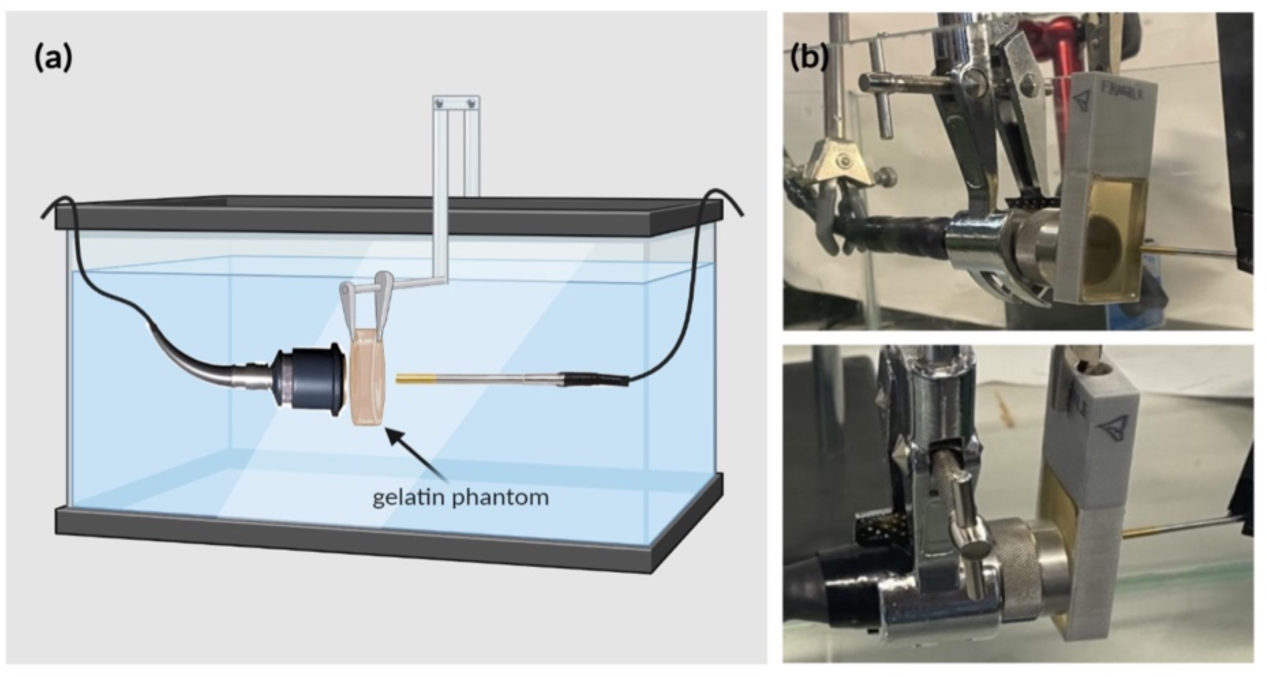
Experimental setup for estimating the acoustic intensity and attenuation coefficient when the gelatin phantoms, of different concentrations, were introduced. (a) schematic diagram of the experimental setup including the gelatin phantom in place in front of the face of the transducer. (b) real-time images showing the ultrasonic signal acquisition setup arrangement, from different views, in which the phantom was held (by the rectangular phantom holder) between the transducer and the hydrophone.

Schematic in Figure 1 describes the acoustic setup built for obtaining the acoustic profiles and the acoustic characterization of the gelatin-based phantoms with all the required connections. The pulsed ultrasonic waves were generated from the 2-channel Siglent arbitrary waveform generator (SDG 2042X; maximum output = 40MHz; maximum sampling rate = 1.2GSa/s; 50 Ω impedance) to which a 50-dB-gain radio frequency (RF) power amplifier was connected (Electronics and Innovation, Rochester New York 14623; 1.0 V_rms_ input; 50 Ω input/output impedance). RF signals from the generator were amplified by the RF power amplifier, before being transmitted through a single element, focused ultrasonic transducer. In general, three focused transducers were used in the experiments with their properties provided in Table 1. The outer diameter of the transducers is 19.05 mm and can be submerged in degassed water for up to 8 hours, long enough for the experiment to be conducted, followed by a dry time of 16 hours to ensure a proper lifetime of the unit. Furthermore, an immersible needle hydrophone (Onda HNR- 0500) was controlled by a 3D-axis motorized system programmed with a LabView sub VI to scan a predetermined volume, then quantize and convert the sound pressure waves emitted by the transducer while traveling through the water tank into voltage signals by the piezoelectric effect. The hydrophone has an operating frequency range of 0.25 MHz to 10 MHz, a maximum operating temperature of 50°C, a capacitance of 200 pF, and an end-of-cable nominal sensitivity of 0.126 V/MPa. Via the data acquisition board (Elvis III, National Instruments; input voltage ≤ 50 V DC, or 30 V_rms_) connected to the hydrophone, the ultrasonic pressure signals were stored as voltage signals, and measured over lateral and longitudinal planes of the ultrasound focal region.

**Table 1.** Properties of the focused transducers used in the experiments

| Manufacturer | Frequency (MHz) | Focal length (mm) | Center Frequency (MHz) | Peak Frequency (MHz) | (-6) dB bandwidth (%) |
| --- | --- | --- | --- | --- | --- |
| Olympus | 1.00 | 37.19 | 0.96 | 1.04 | 73.35 |
| Olympus | 5.00 | 39.52 | 4.77 | 4.86 | 66.52 |
| Olympus | 7.50 | 36.39 | 6.76 | 7.26 | 83.95 |

The acquired voltage signals were processed and analyzed offline through an in-house MATLAB code, and the spatial-peak pulse-average intensity (*I_SPPA_*) was evaluated based on equations *(1)* to *(5)* presented in section “Acoustic measurements”. The ultrasonic voltage data were acquired in water only (Figure 1), and with the phantom placed in between the transducer and the hydrophone (Figure 2). The attenuation coefficient (experimental value) of the phantom was thereafter estimated from these data sets using equation *(S5)*.

For the mechanical assessment of the Young’s modulus of the gelatin-based phantoms, compression tests, using the Universal Testing Machine (825 University Ave, Norwood, MA, US), were conducted on cylindrical specimens of various gelatin concentrations to calculate the mean modulus of elasticity. Each sample of the cylindrical phantom of a particular concentration was placed on a 10-N load cell to be compressed. The aim was to study the effect of increasing phantom gelatin concentration on the elasticity modulus mimicking certain soft biological tissues.

### Phantom Recipe and modeling

The gelatin-based phantoms were prepared using commercially available, low cost, plant-based gelatin. First, 10 mL of osmosed water was preheated to 60°C in a beaker placed on a hot plate. The gelatin powder was then gradually added to the beaker (in the amounts depending on the gelatin concentration needed) and constantly stirred with a magnetic stirrer until totally dissolved. Air bubbles were removed using a fine spatula to eliminate undesirable experimental distortions [24]. The mixture was then poured into a 3D-printed rectangular mold (40 mm by 25 mm) that was previously lightly coated with oil to help in the demolding process following refrigeration. The mold was then transferred to a refrigerator for 24 hours to obtain firm-enough phantoms that can withstand long hours of being submerged in water. Note that the mold was preserved in a covered petri dish to prevent dehydration of the surface. In order to investigate the effect of gelatin concentration on the acoustic attenuation, different phantoms of respective gelatin concentrations (12.5%, 20%, and 24%) were prepared, where an *x* % corresponds to *x* grams of gelatin powder poured in 100 *mL* of water. Following the 24 hours of refrigeration, the phantoms were removed from the fridge, demolded, and mounted in our setup (Figure 2).

### Mechanical Measurements

To test the mechanical properties of phantoms with increasing gelatin concentrations, samples of respective gelatin concentrations of 12.5%, 20% and 24% were prepared. When removed from the fridge and demolded, ten cylindrical specimens of an average height of 8 mm were directly cut from each gelatin-based phantom (of a particular concentration) for mechanical testing. Compression tests were performed on each cylindrical specimen at a displacement rate of 0.1 *mm*/*s* up to a maximum strain of 0.4 *mm*/*mm* using the Instron machine. At room temperature, each specimen was placed on the compression plate with a preload of 0.1 N set for the upper plate to precisely get in contact with the specimen. The stress and strain of each specimen were estimated using the testing machine’s built-in software (Bluehill Universal 3.0), and the mean Young’s modulus was calculated and plotted as a function of gelatin concentration.

### Acoustic Measurements

The motorized setup used LabView VI for system control and data acquisition. The acquired voltage measurements underwent filtering to clean the signal from any noise, and the acoustic intensity of ultrasonic waves was calculated from the filtered voltage matrix according to the guidelines of the Food and Drug Administration [25]. For the computations, the following pipeline was followed: First, the pulse intensity integral (PII) was computed, defined as the time integral of instantaneous intensity integrated over the time in which the hydrophone signal for the specific pulse was nonzero. The instantaneous acoustic intensity *i* is expressed as:

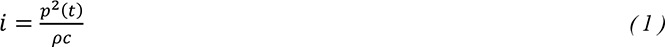

where *p(t)* is the instantaneous acoustic pressure, *ρ* is the density of the medium, and *c* the speed of sound in medium (all in SI units). The pressure *p* is expressed as:

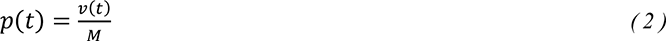

where *v*(*t*) is the acquired voltage array that varies with the position of the hydrophone with respect to the transducer, and *M* is the end-of-cable nominal sensitivity of the hydrophone. Thus, PII is expressed as:

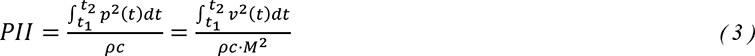

where *(t_2_ – t_1_)* is the interval in which the amplitude of the recorded voltage is nonzero (i.e., t_2_ – t_1_= tone-burst-duration or TBD). PII corresponds to the energy transferred per unit area (expressed in J/m^2^) or the energy fluence during one pulse. Following that, the pulse duration (PD) was calculated, defined as:

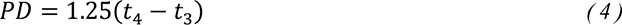

*t_4_* is the time instant at which the time integral of intensity reaches 90% of PII, and *t_3_* is the instant at which the time integral reaches 10% of PII. Finally, the spatial-peak pulse-average intensity (I_SPPA_) was calculated as the maximum ratio of PII (energy fluence per pulse) to PD:

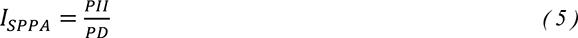

Thus, calculations of the true *I_SPPA_* were not obstructed. With the 1MHz, 5MHz, and 7.5MHz transducers, three different voltages (110 mV, 120 mV and 165 mV) were used to characterize the acoustic profile of ultrasound waves propagating in degassed water, followed by characterizing the effect of gelatin concentration on the acoustic profile with fixed ultrasound frequency and received voltage.

### Acoustic Simulations

Simulations were performed using K-wave, an open-source acoustics MATLAB toolbox, used for modeling the propagation of acoustic waves in well-defined media [26]. The dynamic changes that an acoustic wave undergoes while passing through a 2D compressible medium are expressed with partial differential equations, solved using a k- space pseudo-spectral method. Computations were performed in a 64 by 128 computational grid with a step size of 0.3 mm, enveloped with a perfectly matched layer, occupying 20 grid points covering the circumference of the domain to prevent reflection and wave wrapping. The ultrasound transducer was modeled as a spherically focused acoustic source driven at a frequency of 1MHz and emitting acoustic waves that were focused on a definitive distance of 36 mm from the face of the transducer.

Simulations were carried out in four different settings: acoustic waves propagating in a homogeneous medium containing degassed water only, and heterogeneous media where a gelatin phantom, of three different gelatin concentrations (12.5%, 20% and 24%), was positioned in direct contact with the face of the transducer. Medium properties (sound speed, density, and acoustic absorption coefficient) defining the simulations of water and phantoms of different gelatin concentrations are described in Table S1, such that values were obtained based on the study by Chen et al. [18]. A sensor mask was designed as in Figure S1 to record the acoustic field at each timestep while the acoustic waves propagate through the gelatin phantom and water respectively, taking into consideration the medium properties as they were systematically defined in a piecewise fashion according to the phantom geometry. Calculations of the attenuation coefficient α are detailed in the supplementary based on the studies of [27, 28].

## Results

### Experimental Acoustic Measurements

#### Effect of Frequency, Voltage, and duty cycle on Acoustic Profile and Intensity

To characterize the propagation of ultrasonic waves in water, three transducers were used. Whereby the variation of the acoustic profile and the variation of intensity distribution across the horizontal midplane and vertical transverse sections of the ultrasound beam were studied with the 1 MHz, 5 MHz and 7.5 MHz transducers while increasing the applied voltage (110 mV, 120 mV and 165 mV), and keeping a constant duty cycle of 100 cycles per pulse (Figures 4,5,6). As the ultrasound beams propagated in water, in the x direction, to reach the maximal intensity at the focal point, the size of the horizontal and vertical focal areas increased systematically as a function of increasing applied voltage yet dropped as a function of increased ultrasonic frequency (Figure 8a,b). Note that the horizontal area corresponds to the area of the horizontal midplane, and the vertical area measures the area at the vertical cross section near the focal point and at the focal point, with a cutoff threshold of 85% (inclusive) of the computed maximum intensity. The wider focal area was recorded using the 1 MHz transducer at the highest voltage applied of 165 mV. The higher the voltage and frequency used, the higher the maximal intensity at the focal point reached (Figure 8c).

**Figure 3.**
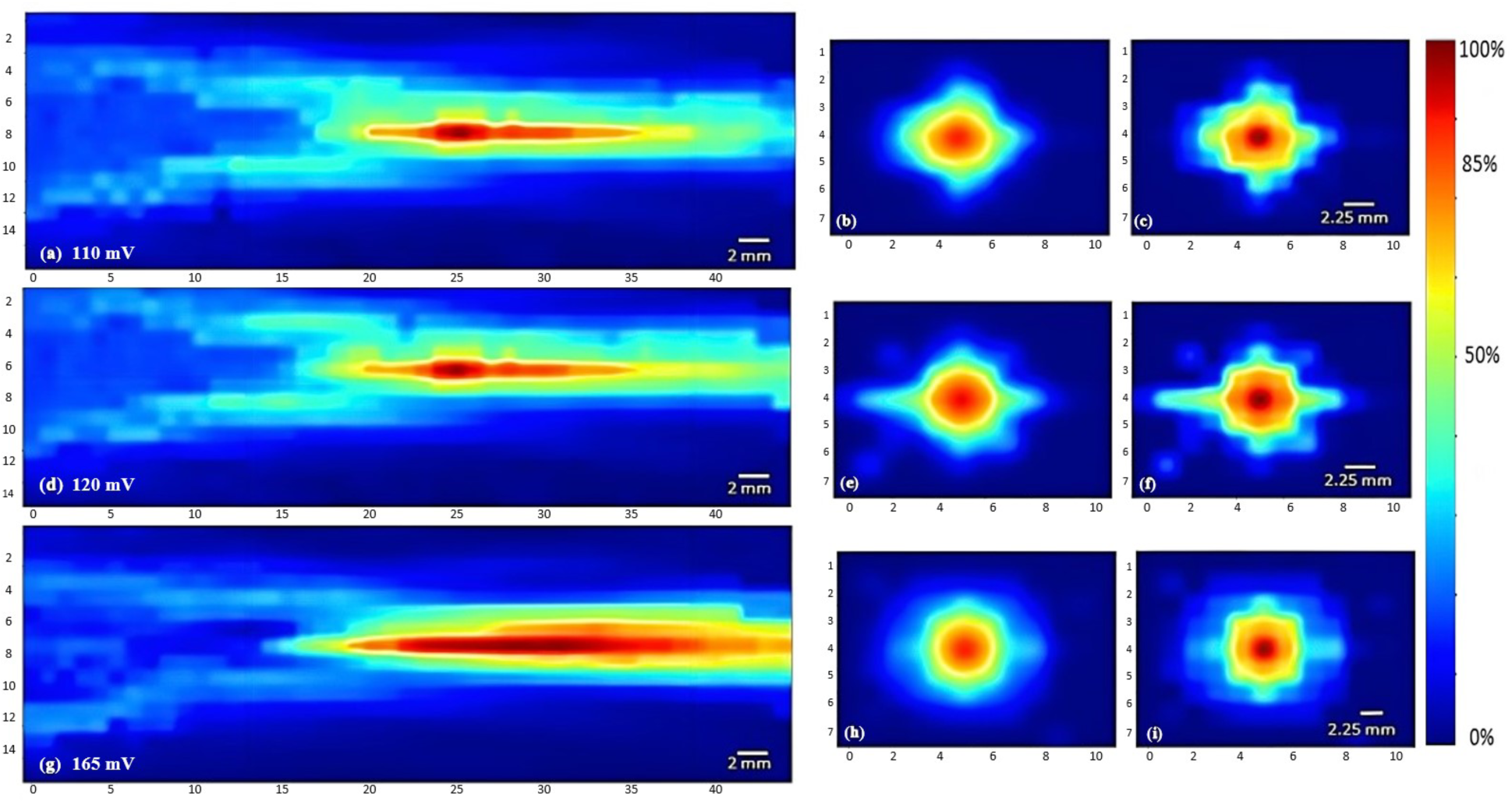
Heat maps of the intensity distribution using the 1 MHz transducer. The horizontal midplane, vertical plane - transverse section near the focal point and at the focal point plotted respectively with a voltage of 110 mV (a, b, c), a voltage of 120 mV (d, e, f), and a voltage of 165 mV (g, h, i).

**Figure 4.**
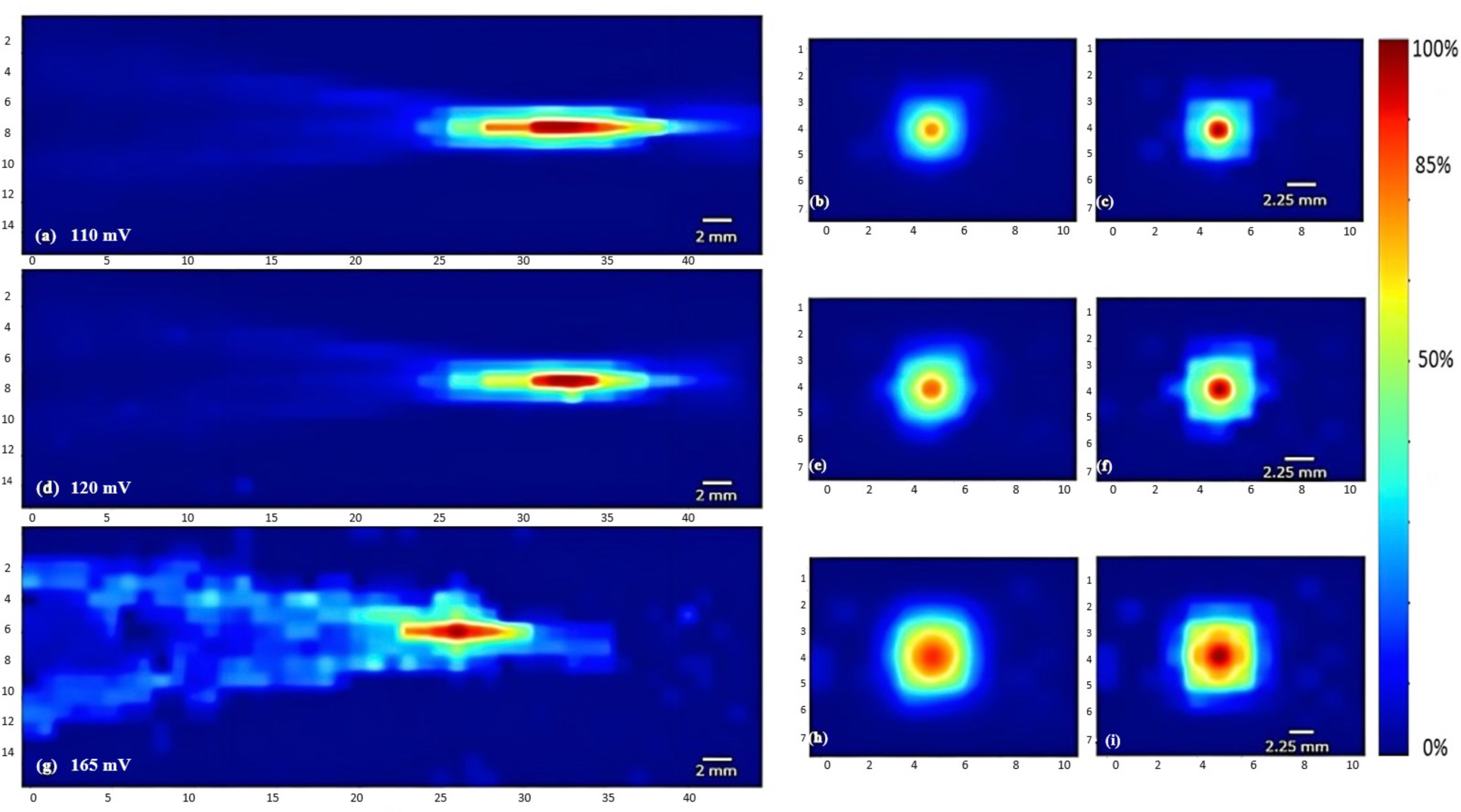
Heat maps of the intensity distribution using the 5 MHz transducer. The horizontal midplane, vertical plane - transverse section near the focal point and at the focal point plotted respectively with a voltage of 110 mV (a, b, c), a voltage of 120 mV (d, e, f), and a voltage of 165 mV (g, h, i).

**Figure 5.**
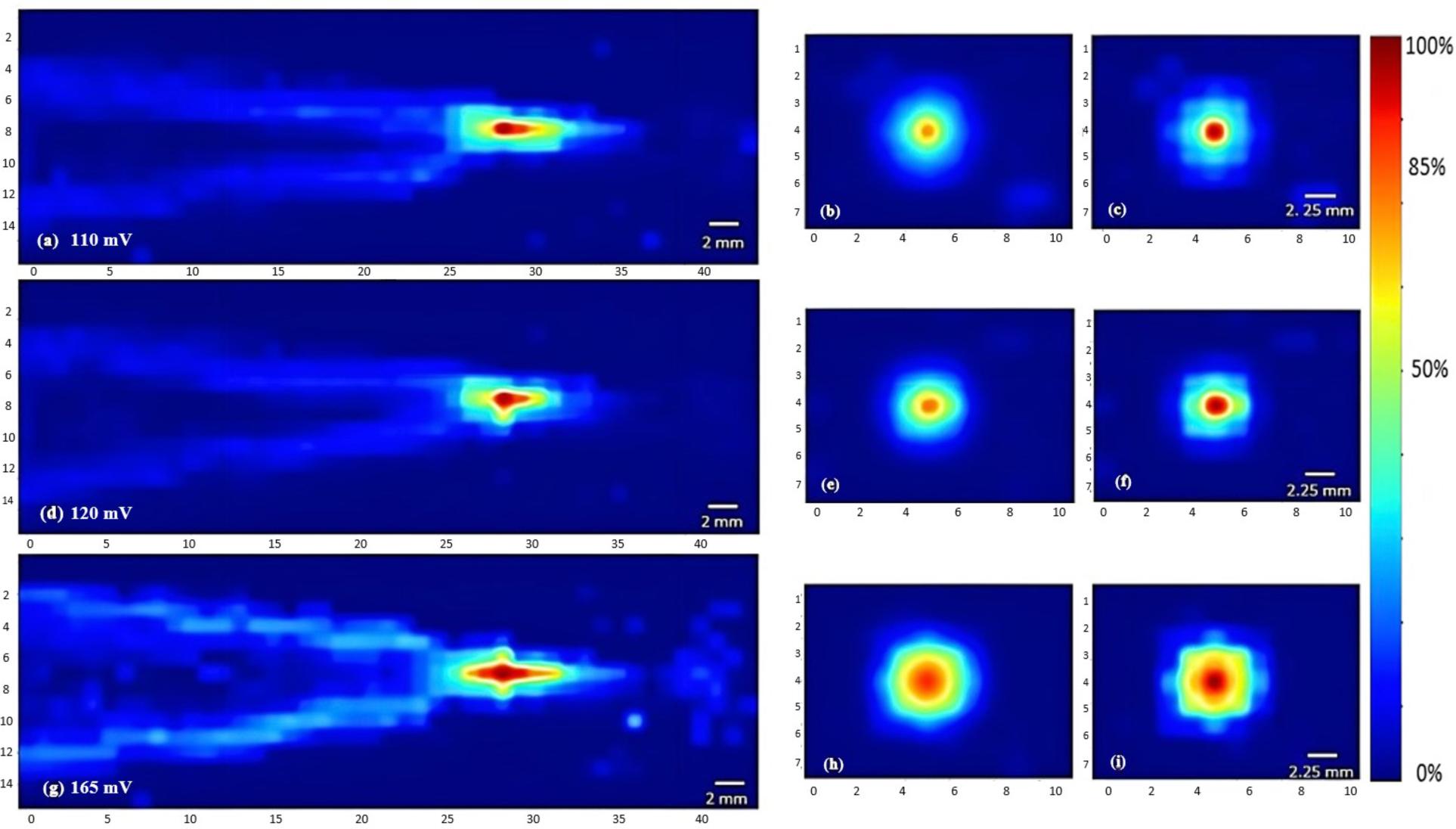
Heat maps of the intensity distribution using the 7.5 MHz transducer. The horizontal midplane, vertical plane - transverse section near the focal point and at the focal point plotted respectively with a voltage of 110 mV (a, b, c), a voltage of 120 mV (d, e, f), and a voltage of 165 mV (g, h, i).

Regarding the effect of varying the duty cycle on the acoustic profile in general and the maximal intensity at the focal point, the 5 MHz transducer was used while receiving a voltage of 120 mV. Incrementing the duty cycle from 33 % to 44 %, followed by 67 %, caused an increase in the maximal intensity (Figure 7d), yet the acoustic profile showed similar intensity spread with all duty cycles used (Figure 6), whereby the horizontal and vertical areas of the focal region demonstrated insignificant change.

**Figure 6.**
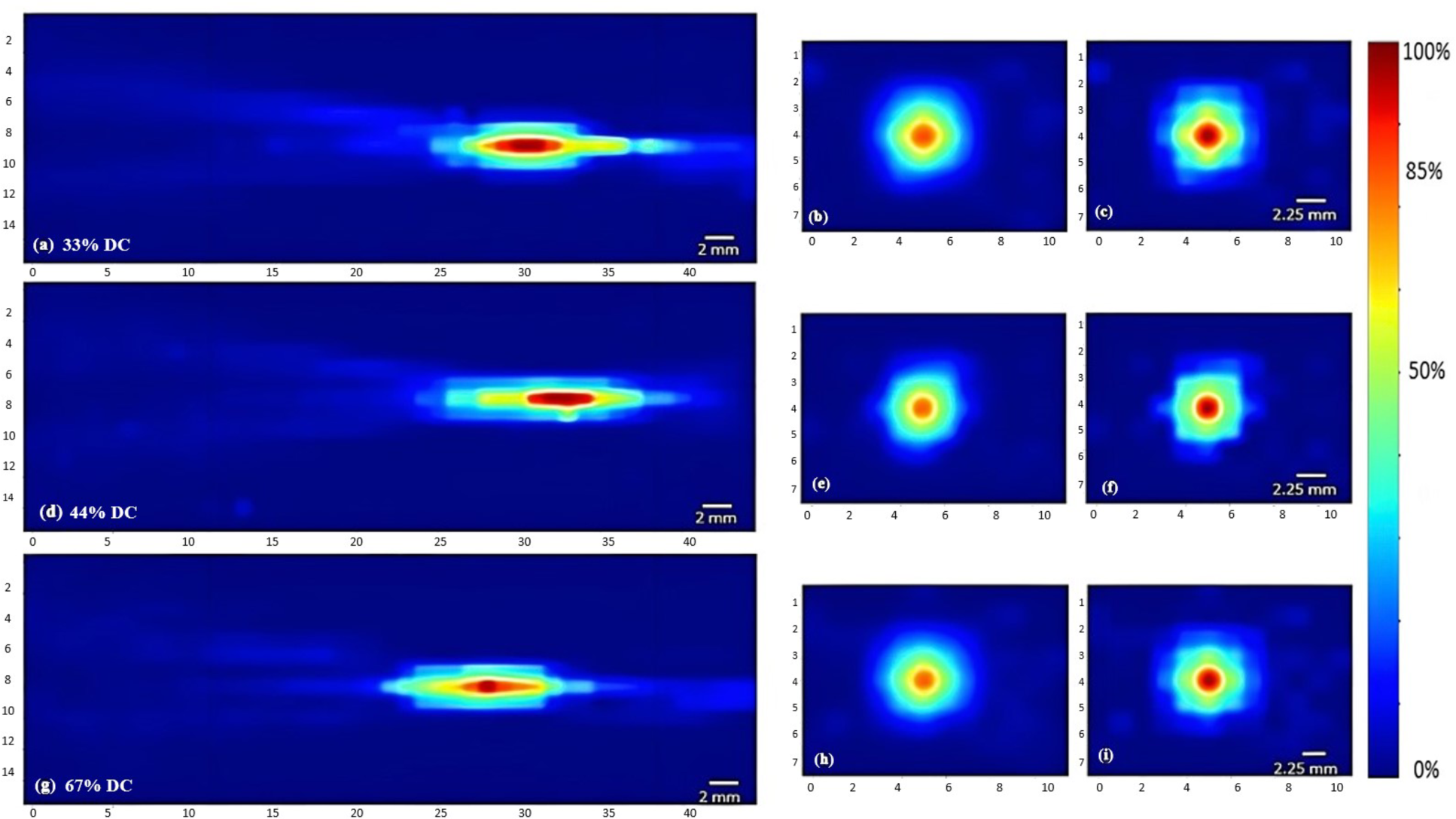
Heat maps of the intensity distribution using the 5 MHz transducer at 120 mV with varying duty cycles. The horizontal midplane, vertical plane - transverse section near the focal point and at the focal point plotted respectively with a duty cycle of 33 % (a, b, c), a duty cycle of 44 % (d, e, f), and a duty cycle of 67 % (g, h, i).

**Figure 7.**
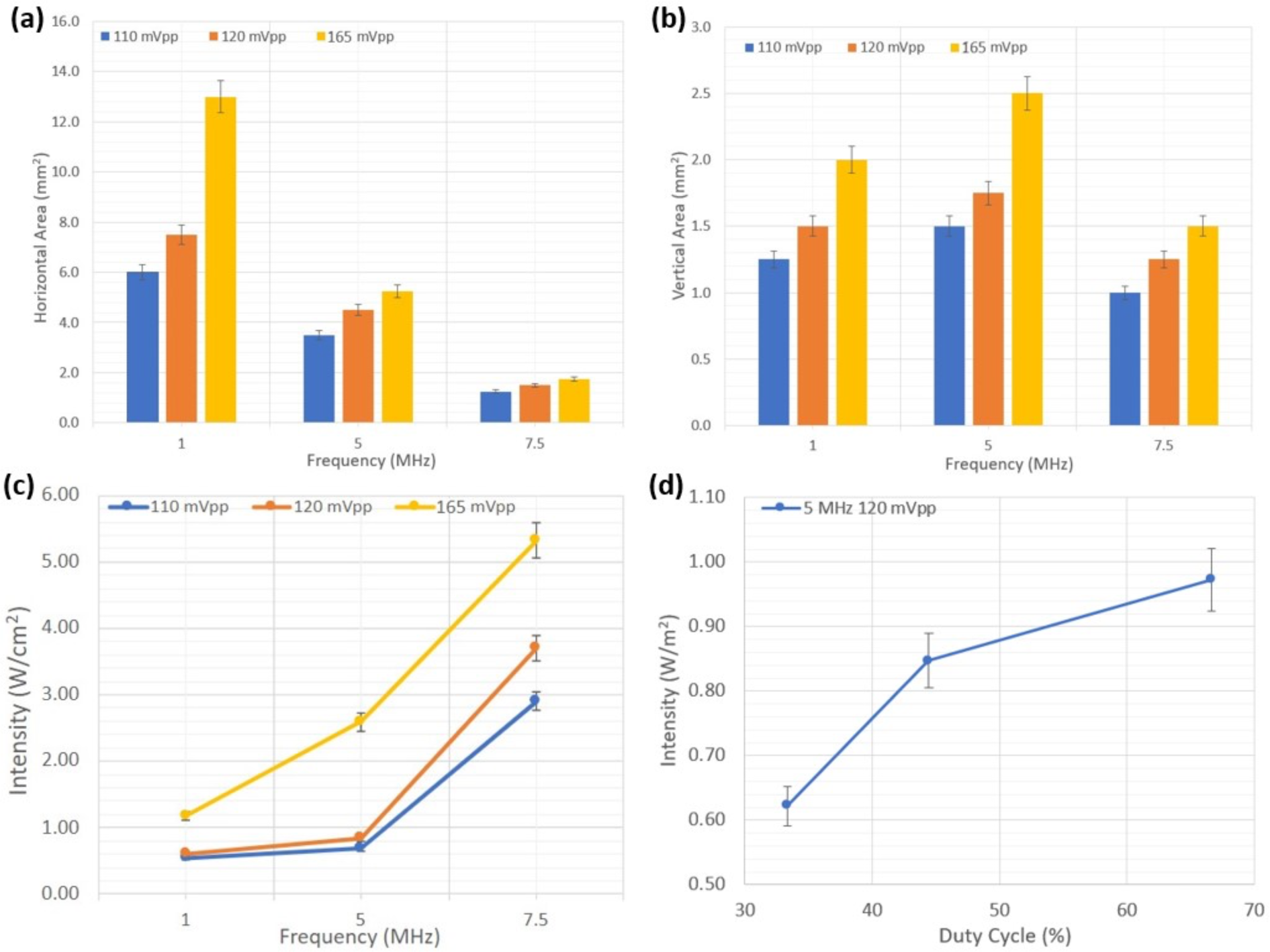
Effect of frequency and duty cycle on the focal area and acoustic intensity. (a) Variation of the horizontal area at the horizontal midplane as a function of ultrasound frequency and voltage. (b) Variation of the area at the vertical cross section at the focal point as a function of ultrasound frequency and voltage. (c) Increase in the maximal intensity at the focal region as a function of increasing acoustic frequency and voltage. (d) Increase in the maximal intensity at the focal region as a function of increased duty cycle with the 5 MHz transducer at a voltage of 120 mV.

#### Acoustic Assessment of Gelatin-based Phantoms

Experimental acoustic characterization of the gelatin-based phantoms was done using our experimental acoustic setup. That same horizontal midplane in front of the face of the transducer was scanned, where all the gelatin-based phantoms of the three different concentrations were placed consecutively. In the first set of experiments, the function generator emitted a signal of 1 MHz fundamental frequency, a burst period of 125 µs (a duty cycle of 80 %), and amplitude of 150 mVpp. The second set of experiments included the 5 MHz transducer, emitting at a burst period of 40 µs and amplitude of 90 mVpp. The maximum *I_SPPA_* was computed from the voltage measurements in each experiment and the attenuation coefficient α was calculated using equation *(S5)*.

**Figure 8.**
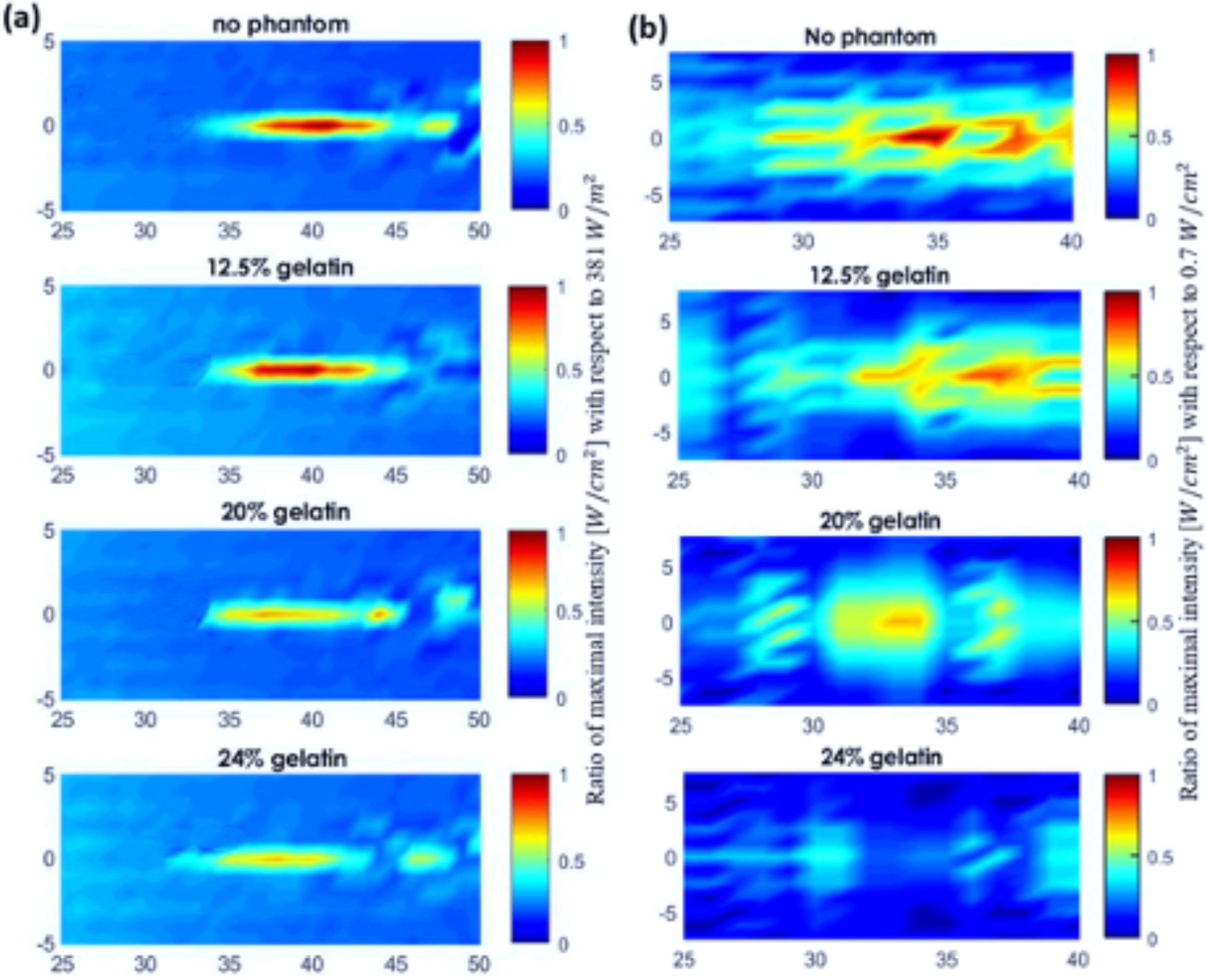
Acoustic profile of the mid-horizontal plane in front of the face of the transducer. (a) The scanned volume without a phantom placed and when the 12.5%, 20%, and 24% gelatin-based phantoms were positioned in place respectively using the 1 MHz transducer (a) and using the 5 MHz transducer (b). The acoustic profiles display the maximal intensities plotted as a function of the maximal intensity recorded in water for visual inspection of the drop in intensity.

Adding more gelatin powder to the solution rendered the gelatin-based phantom more acoustically attenuating where the ultrasound waves emerging from the phantom transferred less power per unit area. The drop in the maximum intensity and subsequent rise in the attenuation coefficient following the increase in gelatin concentration can be visualized in the acoustic scans of Figure 9. The higher the fundamental frequency, the more confined the acoustic profile spread. With the 5 MHz transducer, the focal area, in both the vertical sense and horizontal sense, was narrower than that with the 1 MHz transducer.

**Figure 9.**
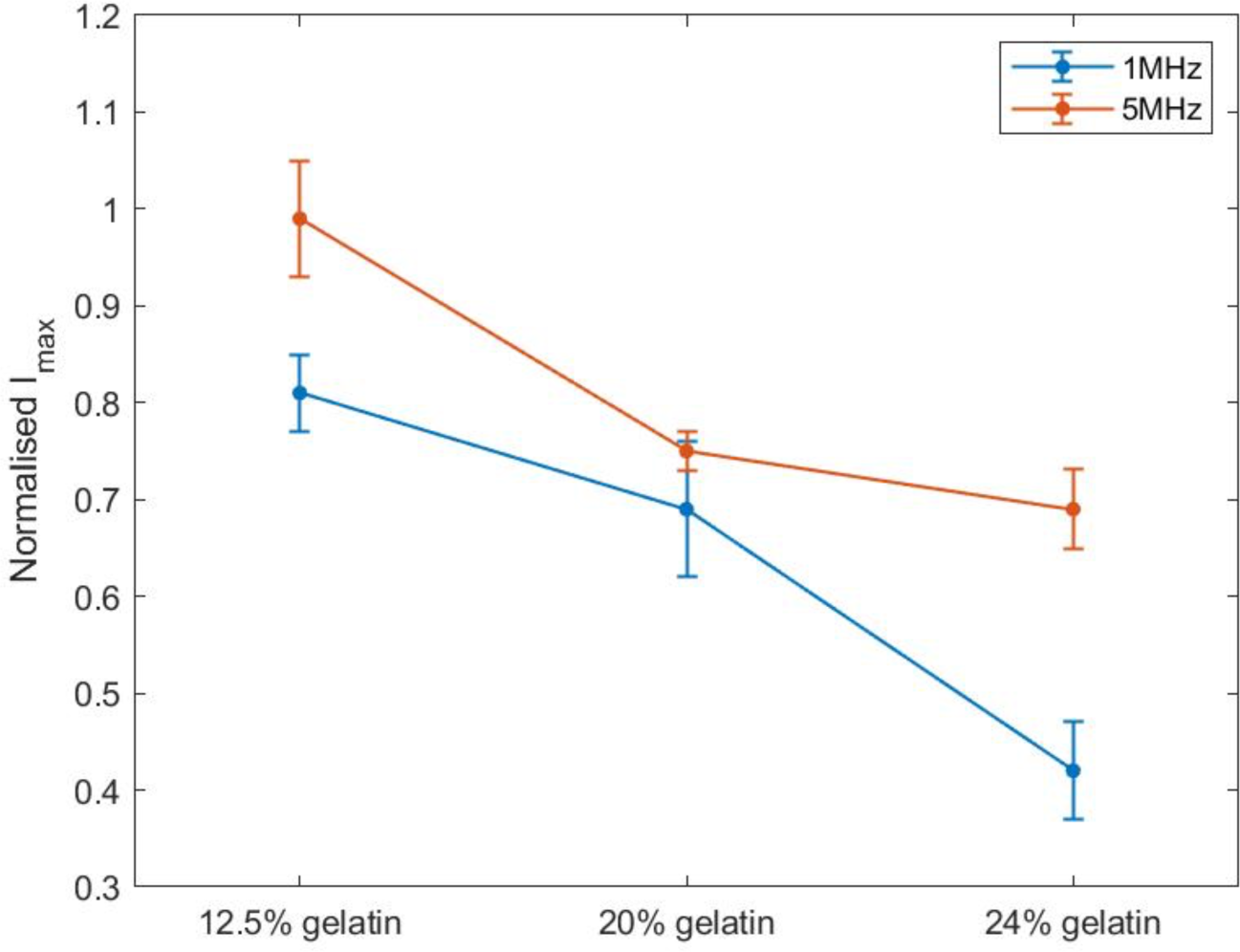
Variation of the normalized intensity with respect to the maximal intensity recorded while using the 1 MHz frequency and 5 MHz frequency transducers.

Computing the normalized intensity with respect to the maximal intensity recorded at the focal point in both experiments showed a decreasing trend with an increase in the gelatin concentration (Figure 10). Add to that, the 5 MHz transducer recorded higher normalized acoustic intensities at all gelatin concentrations when compared with those of the 1 MHz transducer.

**Figure 10.**
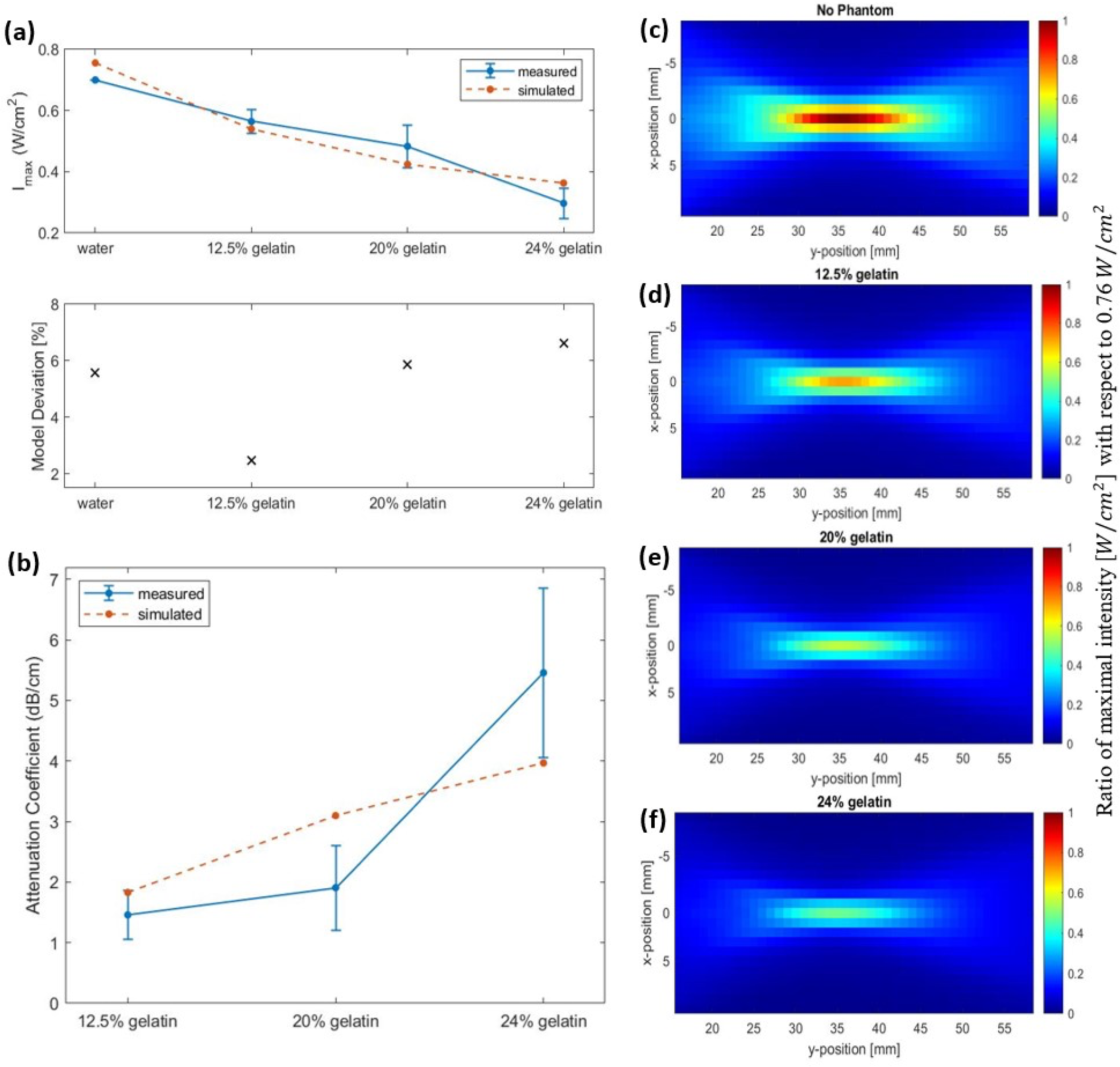
Simulation of acoustic wave propagation through different media. (a) Decrease in the simulated maximum intensity (dashed line) recorded at the focal point in water and as the gelatin concentration increased in comparison with the measured experimental maximal intensity (full line) while using the 1 MHz transducer. The deviation of simulated data from the measured data is less than 10%. (b) Increase in the attenuation coefficient *α* in the simulated (dashed line) and measured experimental data (full line) as a function of increased gelatin concentration. (c) Acoustic profile through water only with a maximum intensity of 0.76 W/cm^3^ reached. (d to f) Visual representation of the acoustic profile where the phantoms with different gelatin concentrations (12.5%, 20% and 24%) were introduced respectively in front of the face of the 1 MHz transducer, showing the decrease in the maximal intensity recorded with respect to that calculated in the water-only medium.

#### Acoustic Profile Simulations

We modeled the propagation of ultrasound waves in the media mimicking biological tissues for effective planning and application. The medium properties and geometries determined the acoustic nonlinearity and power absorption for more accurate modeling. Simulations aided in calculating the maximal intensity at the focal point in front of the face of the 1 MHz transducer which approximately compared to the acoustic fields calculated experimentally. The differences between measured and simulated acoustic fields could arise from various experimental changes including equipment properties, signal detection through the mounted hydrophone and signal processing [29]. Add to that, the phantom properties from dimensions and thickness to refrigeration time and density, affected the attenuation coefficient measurements as well. Nevertheless, the deviation was within an acceptable range for accurate predictions of percentage deviation less than 10% (Figure 10a).

In the simulation experiments, the maximum intensity recorded at the focal point dropped as the gelatin concentration increased (Figure 10a). Simulations showed a similar trend as the experimental data, emphasizing the effect of increasing gelatin concentration on the drop of maximal intensity at the focal point. As a result, the higher the gelatin concentration, the higher the acoustic attenuation (Figure 10b). The effect of increasing gelatin concentration could also be detected in the full volume scans of the acoustic profile in the four settings (Figures 10c–f) where the d, e and f plots illustrated the drop in the maximum intensity recorded when the different gelatin phantoms were introduced with respect to the maximal intensity of 0.76 W/cm^3^ recorded in water only.

#### Experimental Mechanical Measurements

For optimal choice of acoustic parameters, it is crucial to estimate the effect of biological tissues on the intensities of ultrasound waves travelling through them. Consequently, gelatin-based phantoms were prepared and characterized to determine their mechanical properties, particularly Young’s modulus, to be a close mechanical representative of skin and soft-tissues. The Young’s modulus of soft tissues is rather low (in the kilopascal range) when examined at the macroscale of mass tissue assemblies, as being governed by the elastic stiffness of the extracellular matrix proteins [30]. The stress-strain relation for different phantom concentrations is shown in Figure 11a, which demonstrated linearity up to 0.25 mm/mm strain. While increasing the compressive strain, the compressive stress increased, reaching higher values with higher gelatin concentrations. Subsequently, this led to a linear increase in the mean Young’s modulus with the increase in gelatin concentration (Figure 11b).

**Figure 11.**
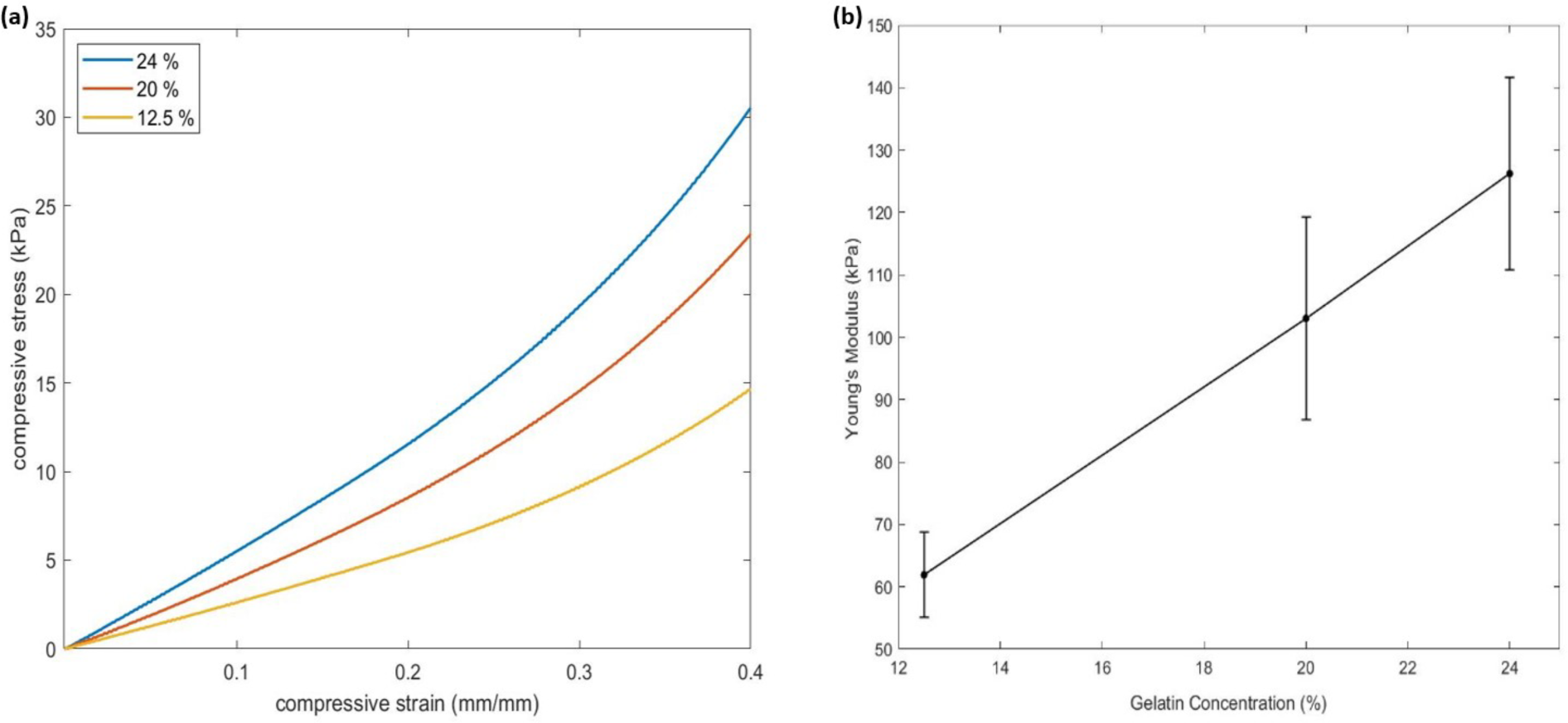
Mechanical properties of the gelatin-based phantoms. (a) Linear stress-strain relation up to 0.25 mm/mm strain of the three gelatin concentrations tested. (b) Increase in Young’s modulus as a function of increasing gelatin concentration where 10 specimens of each concentration were tested

## Discussion

In this work, we utilized high-resolution acoustic mapping to plot the attenuation of ultrasound waves - at different frequencies, voltages, and duty cycles - as they travel through degassed water and soft tissue mimicking gelatin phantoms. Acoustic assessment was performed by recording the intensity drop and lost power per unit area at the focal point of the three transducers and throughout the whole horizontal midplane scanned. For enhanced understanding of the acoustic attenuation of ultrasound waves propagating through the tissue mimicking phantoms, we modeled the acoustic setup and run simulations for outcome prediction and result validation. Whereas the mechanical assessment of the phantoms was estimated by measuring the variation in the Young’s modulus as a function of increasing gelatin concentrations. The compressive stress and strain were measured to study the elastic properties of the gelatin-based phantoms.

### Acoustic Profile Assessment

To study the acoustic profiles of ultrasound transducers with different carrier frequencies, we designed a synchronized semi-automated system that scanned the volume in which ultrasound waves propagated, then processed the data in the form of heatmaps. The focal region of focused ultrasonic transducers was directly related to the operating frequency of the probe, showing a narrowing of the acoustic beam and focal region with the rise in carrier frequency [31]. Extending the “on time” of ultrasound delivery by increasing the duty cycle recorded higher values of acoustic intensities and similar acoustic profiles. For a more confined acoustic profile spread and a higher acoustic intensity reached at the focal point, the 7.5 MHz transducer with a voltage range of 165 mV and ultrasound duty cycle of 67 % achieved the desired results. However, higher frequencies can cause temperature rise when directed towards tissue mimicking phantoms [32]. Decreasing the carrier frequency of the ultrasound transducer allowed more penetrability and a wider acoustic profile spreading across the focal region.

Studying the acoustic properties of the prepared gelatin-based phantom is essential for tissue mimicking in US applications. The acoustic attenuation occurring at the focal point in particular, and the power and energy lost due of mechanical waves in general, can help give a clearer idea of the percentage of intensity drop taking place in tissue. Whether for ultrasound imaging purposes or ultrasound therapies, noninvasive techniques require optimal use of ultrasound parameters to ensure desirable results [33]. The increase in the normalized intensity with respect to the maximal intensity recorded at the focal region when increasing the frequency of ultrasound from 1 MHz to 5 MHz emphasized the results of section 3.1.1 (Figure 9). Whereby, in a given medium with fixed voltage and ultrasound duty cycle, the 5 MHz ultrasound waves transferred more power per unit area. As the gelatin concentration increased, normalized intensities, with both ultrasound frequencies, went down due to less power transfer in a given unit of area as the molds got denser.

Furthermore, we designed a model to generate simulated data and compare it with our experimental results for prediction and validation purposes. The model was used to examine the propagation of acoustic waves as they passed through the gelatin medium to the water medium. Following the variation in the speed of sound and medium density as in Table 1, the amplitude of the received signal at the focal point decreased and the spread of energy was more confined. Gu et al. reported a drop in the amplitude and phase difference between the simulated homogeneous and heterogenous media, mainly from the variation in sound speed, density, absorption, and non-linearity coefficients [34].

The acoustic field predictions and maximal intensity drop at the focal point with respect to those recorded when the different phantoms were placed, one after the other, showed accurate estimations with an acceptable percentage of model deviation (Figure 10a). The variation in the attenuation coefficient calculated between simulated and measured data highly depended on the thickness of the sample (Figure 10b). In the simulation experiments, a fixed thickness of 8 mm was used, while in the experiments of ultrasound waves travelling according to the transmission method, a slight difference in the thickness of the used phantom specimens was recorded. The volume scans including the phantoms clearly illustrated the reduction in the transfer of power per unit area, hence the drop in the intensity values throughout the horizontal plane. The structure, viscosity and concentration of the gelatin gel affected its stress relaxation which caused higher acoustic attenuation and sensitivity to acoustic waves propagation [35].

### Mechanical Assessment of Phantom

Tissues have varying stiffness and elasticity levels ranging from the order of pascal (such as nerve cells and adipose tissues) to the order of Gigapascal (such as bones and tendons) [36]. The mechanical properties of tissues differ depending on their heterogeneity at the microscale and macroscale, this is a result of their varied dimensions, shapes, and constituents. These properties have been outlined using the linear compressive Young’s modulus for the quantification of the tissue stiffness. In addition, mechanical properties of tissue can be altered due to disease enabling detection and diagnosis. For example, Hu et al. reports a mean Young’s modulus of 66.4 ± 12.1 kPa in healthy thyroid nodules of patients while that of cancerous nodules was significantly lower, thus aiding with thyroid diagnosis [37]. In addition, measurements of the Young’s modulus of breast tissues showed high dependance on the tissue precompression applied and tissue constituents, recording values ranging from few kilopascals with normal tissues to folds higher with carcinomas and fibroglandular tissues [38, 39]. In our work, the gelatin concentrations used in our experiments mimicked the elasticity properties of such biological tissues (Figure 12); whereby the mean Young’s modulus estimated for each of the 12.5% and 20% phantom concentrations mimicked skin on the volar and dorsal forearm [40] taking into consideration the skin thickness measurements and layers at these sites; whereas the 24% phantom concentration mimicked the mechanical properties of a pre-stressed patella tendon [41] and certain skeletal muscles [42].

**Figure 12.**
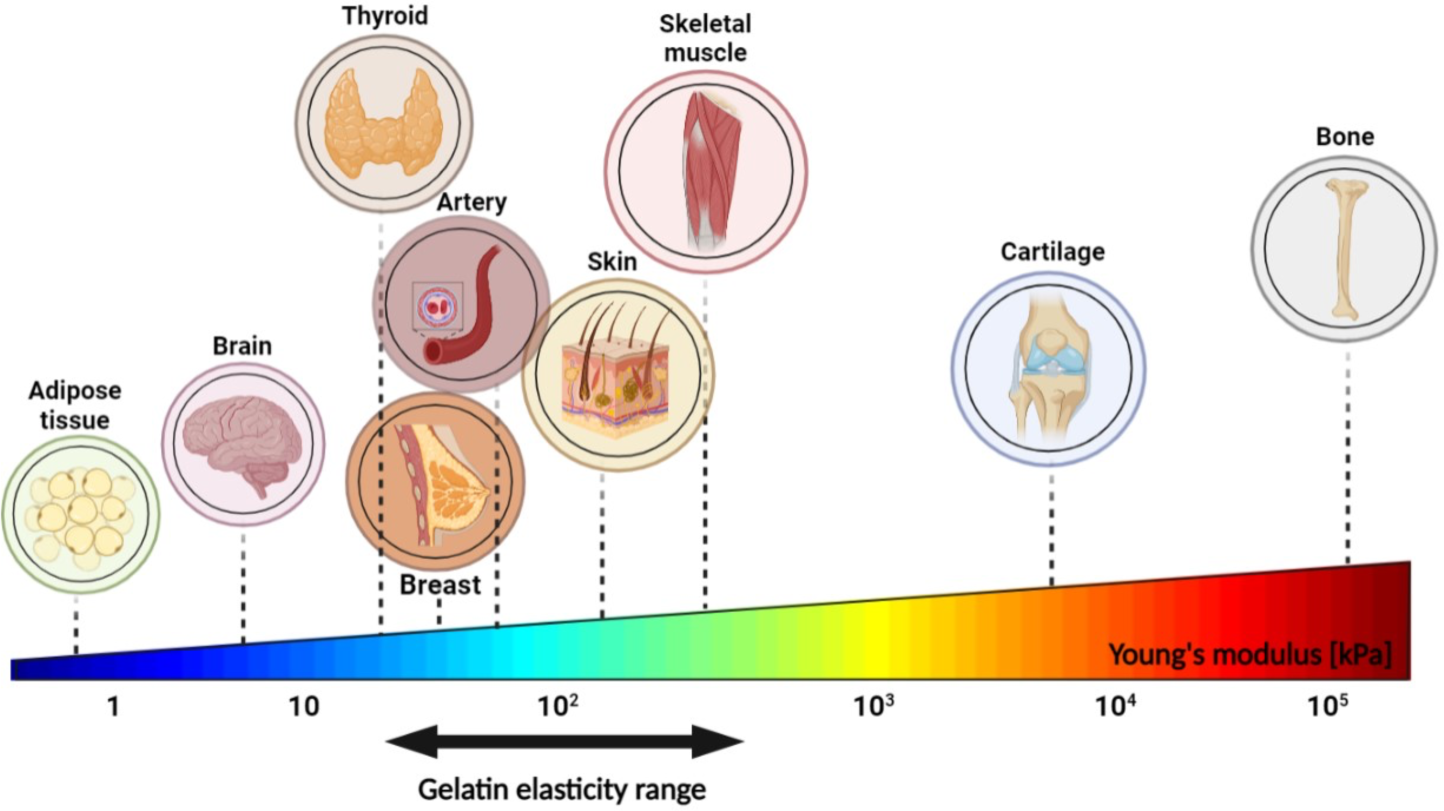
Young’s elasticity modulus of some biological tissues. Gelatin elasticity range of the three different gelatin concentrations used in our mechanical compression experiments mimic the stiffness of some biological tissues.

Many studies document the use of different types of chemicals to prepare mechanically similar tissue mimicking phantoms including water-based phantoms. Polyvinyl alcohol (PVA) cryogel demonstrated adjustable mechanical properties when exposed to freeze-thaw techniques for intravascular elastography [43-45]. Polyacrylamide gels have also been used to fabricate tissue mimicking phantoms with better physical stability than other water-based phantoms [46-48]. In addition, agarose-based phantoms materials are also used in some studies due to their biocompatibility and mechanobiological resemblance of various biological tissues [49]. However, gelatin is regarded as one of the most flexible materials to work with, providing controllable parameters for a rapid and inexpensive recipe for phantom formation. Gelatin records stable elasticity levels and geometry after forming [50]. As for its ultrasonic characterizations, gelatin allows limited scattering levels of ultrasound beams [51]. The range of gelatin percentages used for this study took into consideration the following: a gelatin concentration lower than 12.5% caused the disintegration of the gelatin molds when submerged in water for a long period of time, thus tampering with the integrity of the results. Also, increasing the gelatin concentration beyond 24% risked mixture saturation and heterogeneity [24].

Mechanical testing showed a rise in the elastic modulus with the addition of more gelatin powder to the solution. This could be a result of denser molds having higher stiffness and firmness after refrigeration. The variability in the Young’s moduli of the samples recorded experimentally could be a consequence of slight dehydration, as the sample phantoms were removed from the fridge, demolded and specimens were cut out from them; Kalra et al. reported a significant reduction in Young’s modulus of the outermost skin layer as a function of increased hydration [52], thus affecting the skin’s mechanical properties. It should be noted that specimens obtained from the edges of the prepared phantom molds record higher stiffness values than those cut out from the center, given that the gelatin hardens faster at the surface and boundaries than in the middle deeper areas. Humidity index and room temperature could also increase the degree of discrepancy. In addition, the variability increased as the specimens became denser and less homogeneous thus augmenting the standard deviations. In Fig. 11a, and for the same stress applied, smaller strain was recorded with higher gelatin concentrations; whereby adding more gelatin powder rendered the phantoms denser and more capable of withstanding compressive deformations in the direction of the applied force. Finally, the phantoms had its limitations as it could only withstand being submerged in water for a certain period of time. Extended submergence would cause the phantom to start disintegrating and thinning out at specific regions, thus interfering with the integrity of the results.

## Conclusion

Acoustic phantoms are a valuable tool for researchers in various fields, providing a safe and reliable way to study the behavior of sound waves. They are also used to test the accuracy of acoustic measurements and to evaluate the performance of acoustic devices. In this work, developed gelatin-based ultrasound (US) phantoms of soft tissue, mechanical via unconfined compression tests, acoustically via high resolution acoustic mapping and validated the results against acoustic simulations. High resolution acoustic maps of the intensity distribution of US can provide essential information on the spatial changes in US wave intensity and focal point, enabling a more in-depth examination of the effect of tissue on US waves. Our work, described the acoustic and mechanical characterizations of a phantom suitable for investigating the effective intensity of an ultrasonic wave after encountering a soft tissue. High-resolution acoustic mappings allow visualization of multidimensional measurements of spatio-temporal acoustic waves. Examining the ultrasound intensity drops as a function of increased tissue elasticity and stiffness, mimicked by artificial *in vitro* phantoms, highlighted the acoustic attenuations taking place. As ultrasonic waves pass through different media, the magnitude of energy loss goes up. Estimation of the latter enhances pre-clinical and clinical works that employ ultrasound. Advances in computational simulations of the acoustic profiles facilitate prediction and validation of the experimental outcomes for fitter results. For further enhancements and more precise measurements, the phantoms can be fabricated in more accurate geometrical shapes, specific to the tissue in concern. Henceforth, this work presented a method by which a readily available material can be characterized to mimic properties of soft tissues, using a motorized system to get high resolution acoustic intensities and profiles of the traveling ultrasonic waves in characterized media.

