## Supplementary for "High Resolution Acoustic Mapping of Gelatin-Based Soft Tissue Phantoms"

Propagation of ultrasound waves through the gelatin-based phantoms resulted in acoustic attenuations of the transfer of ultrasound intensity throughout the medium. Whereby the sound pressure wave, of a certain amplitude *P*, experienced acoustic alterations as it traveled through the phantom. Comparing the wave amplitude in water with that through the phantom of thickness $x$, attenuation was calculated based on the following equations:

$$P_{phantom} = P_{water} e^{-\mu x} \quad (S1)$$

where μ is the amplitude attenuation coefficient in Np/cm.

Noting that *I* is directly proportional to $P^2$, this implies:

$$\frac{P_{phantom}}{\sqrt{I_{phantom}}} = \frac{P_{water}}{\sqrt{I_{water}}} \quad (S2)$$

Manipulating equations *(6)* and *(7),* we get $I_{phantom} = I_{water} e^{-2\mu x} \Rightarrow x = -\frac{ln\left(\frac{I_{phantom}}{I_{water}}\right)}{2\mu}$ (S3)

Furthermore, changing the basis of logarithm to base 10 and simplifying equation *(8)* we get,

$$8.686\mu = -\frac{10\ log\left(\frac{I_{phantom}}{I_{water}}\right)[dB]}{x\ [cm]} \quad (S4)$$

By definition, 8.686 μ is the intensity attenuation coefficient α in dB/cm [34]. As simulations were run, the maximum intensity was recorded at the point of focus (i.e., spatial-peak pulse-average intensity) and the attenuation coefficient $\alpha$ of the acoustic waves as they pass through the phantom was calculated according to the following equation:

$$\alpha = -10 \frac{log(\frac{I_{phantom}}{I_{water}})}{x} \quad (S5)$$

where $I_{water}$ and $I_{phantom}$ define the maximum intensity recorded in water and when the different phantoms were immersed respectively. The average thickness $x$ of each phantom was estimated to be 8 mm.

**Table S1.** Medium properties of water and phantoms of different gelatin concentrations used to simulate the media through which the acoustic waves propagate

| | | Sound speed $[m.s^{-1}]$ | Density $[kg.m^{-3}]$ | Absorption coefficient $\alpha_0$ $[dB.cm^{-1}.MHz^{-1}]$ |
|---|---|---|---|---|
| Water | | 1500 | 1000 | 0.0025 |
| Gelatin | 12.5 % | 1545 | 1040 | 0.1275 |
| | 20 % | 1571 | 1063 | 0.204 |
| | 24 % | 1585 | 1080 | 0.245 |

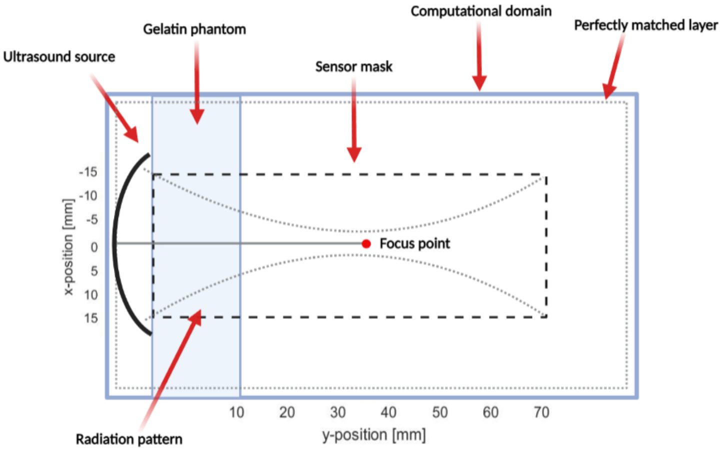


**Fig. S1** Schematic representation (in K-wave) of the computational domain including the position of the spherically focused ultrasound transducer and its focal point, rectangular gelatin phantom and the perfectly matched layer around the domain edges for suppression of wave reflection. The sensor mask was modeled to cover the whole radiation pattern of the ultrasound transducer
